# Harnessing fusion of genome-edited human stem cells to rapidly screen for novel protein functions *in vivo*

**DOI:** 10.1101/2025.06.25.661608

**Authors:** Samantha L. Smith, Yuichiro Iwamoto, Aadhithya Manimaran, David G. Drubin

## Abstract

Genome editing has enabled the integration of fluorescent protein coding sequences into genomes, resulting in expression of in-frame fusion proteins under the control of their natural gene regulatory sequences. While this technique overcomes the well-documented artifacts associated with gene overexpression, editing genomes of metazoan cells incurs a significant time cost compared to simpler organisms, such as yeast. Editing two or more genes to express multiple fluorescent fusion proteins in a single cell line has proven to be a powerful strategy for uncovering spatio-dynamic, and therefore functional, relationships among different proteins, but it can take many months to edit each gene within the same cell line. Here, by utilizing cell fusions, we quickly generated cells expressing pairwise permutations of fluorescent fusion proteins in genome-edited human cells to reveal previously undetected protein-organelle interactions. We fused human induced pluripotent stem cells (hiPSCs) that express in-frame fusions of clathrin-mediated endocytosis (CME) and actin cytoskeleton proteins with hiPSCs that express fluorescently tagged organelle markers, uncovering novel interactions between CME proteins, branched actin filament networks, and lysosomes.

**Significance Statement:**

- Cell fusion can be used to generate new multi-colored genome-edited cell lines, which can be clonally expanded.
- In combination with genome-edited fluorescent cell line libraries such as the collection from the Allen Institute for Cell Science, cell fusion can be used as a screening tool to look for novel protein localization on a variety of intracellular structures.
- Multiple endocytic proteins localize to the surface of lysosomes in healthy, non-stressed cells, and these proteins exhibit a broad range of behaviors on the lysosome surface.

## Introduction

Cell lines engineered to express multiple proteins as fluorescent protein fusions of different colors have proved useful for revealing the intricate spatial and dynamic regulation of cellular processes, ranging from mitosis to motility to membrane trafficking. For such studies, endogenous tagging methods employing genome editing have proven superior to traditional over-expression methods that often result in aberrant protein localization and dynamics (Bukhari & Müller, 2019; Doyon et al., 2011; Gibson et al., 2013; Ratz et al., 2015). However, construction of multi-colored mammalian cell lines requires successive rounds of gene editing, which is very time-consuming, limiting the number of pairwise permutations of tagged protein combinations that can be examined. This is because making even a single-colored mammalian cell line requires designing and cloning a donor plasmid and single-guide RNA construct, performing the editing, fluorescence-activated cell sorting (FACS), clonal expansion, and validation (PCR or whole genome sequencing and Western Blot analysis). Assuming these steps go according to plan, construction of a single fluorophore knock-in cell line can take a month or longer, depending on the cell type. Successive editing is necessary rather than making multiple edits simultaneously because genetic knock-in editing efficiency for a single tag is very low, much lower than for genetic knock-outs, and thus, trying to knock in multiple tags reduces the success rate even further (Leal et al., 2024). Additionally, editing multiple genes simultaneously has the potential for higher off-target events, as when using multiple guide RNAs, there is a higher chance that one might bind to a site with few mismatches, resulting in off-target cleavage, and requires a higher concentration of Cas9 enzyme, also increasing the chance of binding to off-target sites (Chen et al., 2023; Tian et al., 2023; Veluchamy et al., 2023; Zhang et al., 2015). Most importantly, it is essential to demonstrate that each tag results in a functional protein before making additional edits.

As genome editing has become a staple in research, labs and institutions are generating large banks of genome-edited cell lines (Roberts et al., 2017), which are invaluable shared resources. Cell fusion offers the possibility of combinatorially fusing different combinations of cell lines to rapidly investigate new biology in a time-efficient manner. In this study, to circumvent the need to perform multiple rounds of genome editing to generate multi-colored cell lines, we utilized polyethylene glycol-1500 (PEG) to fuse mammalian cells and generate multi-colored cell lines. We show that this approach is effective for standard tissue culture lines as well as for human induced pluripotent stem cell lines (hiPSC). hiPSC lines offer many advantages over tissue culture cells (Drubin & Hyman, 2017). We conducted a systematic screen by surveying different combinations of organelle markers and markers for clathrin-mediated endocytosis proteins using cell fusion. We identified several interactions, demonstrating the power of this approach. This study demonstrates the strength and potential of cell fusion as a screening method to investigate novel protein functions in a time-efficient manner that takes advantage of existing genome-edited alleles present in different cell lines.

## Results

### Generation of dual-tagged SK-MEL-2 cells through fusion of single-tagged, genome-edited cells

To validate our cell fusion method, we began by fusing SK-MEL-2 melanoma cells genome-edited to express clathrin-RFP (Clta-RFP) or dynamin2-GFP (Dnm2-GFP) from their endogenous genomic loci (Doyon et al., 2011). These markers were chosen because of their well-characterized protein dynamics during CME (Aguet et al., 2013; Avinoam et al., 2015; Doyon et al., 2011; Kaplan et al., 2022). Clathrin self-assembles into a cage around the nascent endocytic vesicle (Brodsky, 2016). Dynamin2 is a GTPase responsible for the scission of the vesicle from the plasma membrane for intracellular trafficking (González-Jamett et al., 2013). Together, these two tagged CME protein markers allow the tracking of an endocytic event from the early stage, marked by clathrin recruitment, to the end of the event, marked by dynamin2 recruitment. Following the method outlined in (Agoglia et al., 2021), we fused cells by incubating two cell lines, each expressing only one tagged protein, with PEG. We allowed them to recover for one day before examination. Following PEG incubation, fused cells were larger than non-fused cells when observed by phase-contrast microscopy. Total Internal Reflection Fluorescence (TIRF) microscopy was then used to image Clta-RFP and Dnm2-GFP cells that were either co-cultured in a pooled population before fusion, or pooled and then incubated with PEG. Our results demonstrated that in the absence of PEG, the pooled cells only expressed either Clta-RFP or Dnm2-GFP as punctate structures at the basal surface, while the fused cells expressed both endocytic markers (Figure 1, A and B).

**Figure 1.**
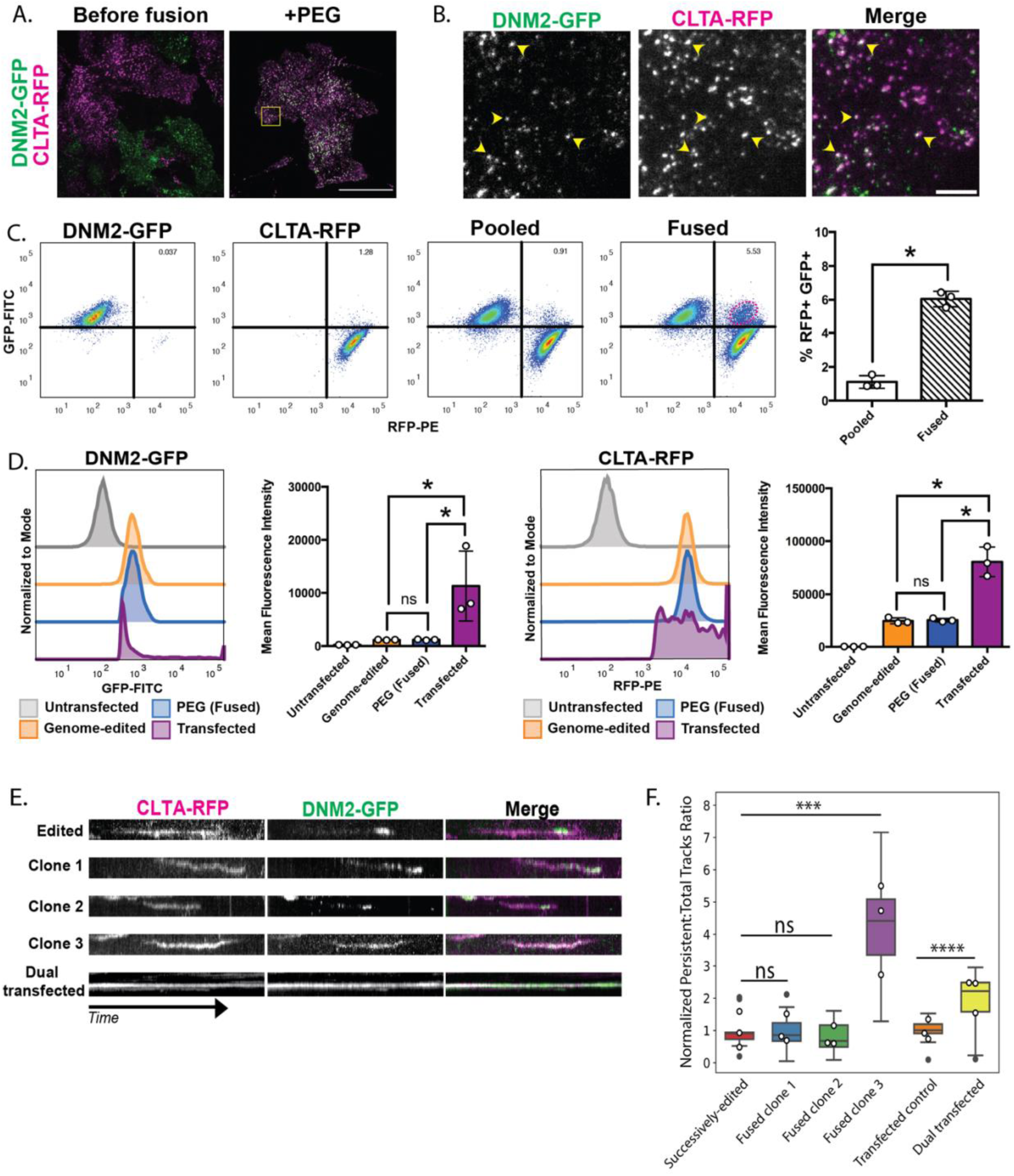
(A) Total Internal Reflection Fluorescence (TIRF) still image showing the two parent cells co-cultured without PEG fusogen where the magenta and green signals reside in different cells (left), and the two parent cells co-cultured with fusogen where the magenta and green signals are now present in the same fused cells (right). Scale bar = 40µm. (B) Inset from the fused cell image in the yellow box in (A) showing co-localization of DNM2-GFP and Clta-RFP (yellow arrowheads). Scale bar = 5 µm. (C) Flow cytometry data from a fusion experiment. The signal from the PE laser (RFP) is on the x-axis, and the signal from the FITC laser (GFP) is on the y-axis. The first plot represents the DNM2-GFP parent cell line, the second plot represents the Clta-RFP parent cell line, the third plot represents the two parent cells co-cultured without fusogen, and the fourth plot represents the two parent cells co-cultured with fusogen. Quadrants are gated to measure the enrichment of a new RFP+ GFP+ population, encircled by a red dotted line. The percentage of RFP+ GFP+ cells in the total population with and without fusogen across three replicates is shown in the bar graph. (D) Histograms representing GFP-FITC and RFP-PE signal intensity for untransfected cells that do not express a GFP or RFP fusion protein, genome-edited cells that express a GFP or RFP fusion protein, PEG-treated (fused) GFP+ RFP+ cells, and cells transfected to overexpress the GFP and RFP fusion proteins. Quantification of the mean fluorescence intensity for both GFP-FITC and RFP-PE across three replicates is shown in the bar graphs. (E) Kymographs of single CME events from three examples of clonally expanded dual-colored cells after fusion, and cells over-expressing clathrin-RFP and dynamin-GFP. Time is on the y-axis (2 minutes). (F) Bar graph plotting the ratio of persistent tracks to total tracks. Fused clones 1, 2 and 3 are normalized to the successively genome-edited control (n = 9, ***, P < 0.05, for Mann-Whitney test), and the dual transfection (CLTA-RFP and DNM2-GFP) is normalized to the transfection, successively genome-edited control (n = 15 for the control and n = 23 for the transfected, ****, P < 0.05, for Mann-Whitney test). The box plot shows the upper and lower quartiles with the median value displayed as a line. The white dots on top of each bar represent the mean ratio per experimental replicate.

To determine the efficiency of cell fusion, we next quantified the percentage of the total population that was fused. To do this, we used flow cytometry to measure the levels of RFP and GFP in individual cells within a population (Figure 1C). We found that there were distinct populations of GFP-positive and RFP-positive cells in the pooled but non-fused population representing the two single-tagged cell lines (Figure 1C). After the cells were incubated in PEG, there was a marked enrichment in the percentage of single cells that were both GFP positive and RFP positive from ∼1% in the pooled condition to ∼6% in the PEG-treated population (Figure 1C). We hypothesize that the cells scored as dual positive in the non-fused, pooled condition might be auto-fluorescing dead cells. From these experiments, we determined that we can successfully fuse cells to generate dual-tagged cells. While the percentage of fused cells is less than 10%, we felt our conditions were optimal because longer PEG incubation resulted in increased cell death. We also used flow cytometry to measure the fluorescence intensity of GFP- and RFP-tagged proteins at the single-cell level to assess any variance in protein expression. For these experiments, we also included cells transiently transfected with constructs to express both Dnm2-GFP and Clta-RFP to see how these proteins behaved in genome-edited, fused cells compared to cells over-expressing the two proteins. At the single cell level, the fused cells show a normal distribution of Dnm2-GFP and Clta-RFP fluorescent signal and mean fluorescence intensity compared to the non-fused, genome-edited control cells (Figure 1D). Importantly, the transfected cells show a very irregular fluorescent signal intensity distribution, with a much higher overall fluorescence intensity, and a large standard deviation of fluorescence intensity compared to both non-fused and fused genome-edited cells (Figure 1, D and F, S1A and B). Under transient transfection conditions, we see a larger standard deviation, which attests to the amount of heterogeneity between cells within the same experiment and suggests that fusion of genome-edited cells is a more reliable method than transient transfection for studying protein dynamics and interactions in live cells.

To investigate the DNA content of fused cells, we also incubated these samples with Hoechst-33342 dye prior to flow cytometry. In the pooled, non-fused population, we observed a peak in the Hoechst fluorescence intensity, corresponding to cells with the expected diploid (2n) DNA content. When applying this same gating to the GFP+ RFP+ cells in the PEG-fused sample, there was a shift toward increased Hoechst signal, indicating that a larger portion of single cells had higher DNA content resulting from fusion, with a small shoulder on the right of the curve representing the population of cells with higher DNA content that most likely were undergoing mitosis (Figure S1C). From these experiments, it was clear that we could successfully fuse cells to generate dual-tagged cells and that the protein expression in the fused cells was more similar to expression in unfused, genome-edited cells than in transfected cells. We next asked whether CME protein dynamics were also conserved in the fused cells.

### Fused, polyploid SK-MEL-2s can be clonally expanded and retain endocytic dynamics characteristic of successively genome-edited parental cells

To determine whether normal endocytic dynamics were preserved in fused cells, we first sought to obtain a more homogeneous population of fused cells. To do this, we used fluorescence-activated cell sorting (FACS) to isolate individual fused cells expressing both Clta-RFP and Dnm2-GFP for clonal expansion. Using the same gating scheme as described above, we observed the enrichment of SK-MEL-2 cells positive for both RFP and GFP signals in the PEG incubated population and formed a gate around this enriched population to use for sorting. We also incubated these samples in Hoechst 33342 DNA dye prior to FACS sorting. We formed a gate around the range of Hoechst fluorescence intensity that was greater than the 2N peak to enrich cells with increased DNA content expected for fused cells. The population contained within both gates was sorted as single cells into a 96-well plate using FACS and clonally expanded.

Three independent clones from the sorting were selected for live-cell imaging. For these studies, we included normal, genome-edited cells expressing both Clta-RFP and Dnm2-GFP and a transiently transfected, non-fused cell line overexpressing both Clta-RFP and Dnm2-GFP. CME dynamics were analyzed in the three cell types. We used kymographs created from 4-minute movies to compare endocytic dynamics in the three cell types qualitatively. We observed that CME dynamics in the fused cells were more similar to those in the genome-edited control, with clathrin internalization coinciding with dynamin recruitment, which marks vesicle scission (Figure 1E, Movie 1). We also used particle tracking software to compare endocytic dynamics in these cell types quantitatively and found that, for sites with lifetimes of 20-180 seconds, CME sites in the fused clones recruit clathrin before dynamin similar to the non-fused, genome-edited cells (Figure S1D). Lastly, we also quantified the ratio of persistent tracks to total tracks. Persistent tracks are classified as sites that last the duration of the 4-minute movie and are more prevalent in transfected cells over-expressing Clta-RFP and Dnm2-GFP compared to genome-edited cells (Figure 1F, Doyon et al., 2011). In two of the three fused clones, there were fewer persistent tracks compared to transiently transfected cells, which supports the conclusion that fused cells display native protein dynamics (Figure 1F). However, a third clone showed a pronounced increase in the ratio of persistent to normal tracks, which highlights the importance of characterizing multiple clones after sorting. Additionally, with PEG fusion being a rapid treatment, it is feasible to investigate protein dynamics across multiple non-clonally expanded cells relatively easily and quickly. Overall, these results indicate that protein dynamics are better preserved in cell lines created by cell fusion than in cell lines created by transient transfection.

### Fusion of genome-edited hiPSCs enables combinatorial imaging of encoded fluorescent proteins

Our results in SK-MEL-2 cancer cell lines led us to wonder if we could employ the same techniques on a non-cancer cell line, so we turned to human induced pluripotent stem cells (hiPSCs). We started by treating two separately genome-edited hiPSC lines, one expressing AP2-TagRFP-T and Dnm2-tagGFP2 and the other AP2-TagRFP-T and ArpC3-HaloTag, with PEG (Figure 2A). AP2 is a heterotetrametric protein complex that is recruited early in the endocytic pathway to cluster cargo molecules at the endocytic site and it bridges the clathrin coat to the plasma membrane (Smythe, 2002). ArpC3 is a component of the Arp2/3 complex, which nucleates assembly of branched actin filaments around endocytic sites to aid in vesicle internalization (Galletta & Cooper, 2009; Rottner et al., 2017). Expression of all of these markers in one three-colored fusion cell line would allow us to monitor coat formation, actin assembly and vesicle scission at individual CME sites. We confirmed that cell fusion was occurring in hiPSCs by staining them with Hoechst DNA dye and a plasma membrane marker, CellMask (Figure S2A). Before fusion, we observed distinct cell boundaries between cells expressing GFP and HaloTag, but after fusion, we observed cells expressing all three fluorescent tags (Figure 2A).

**Figure 2:**
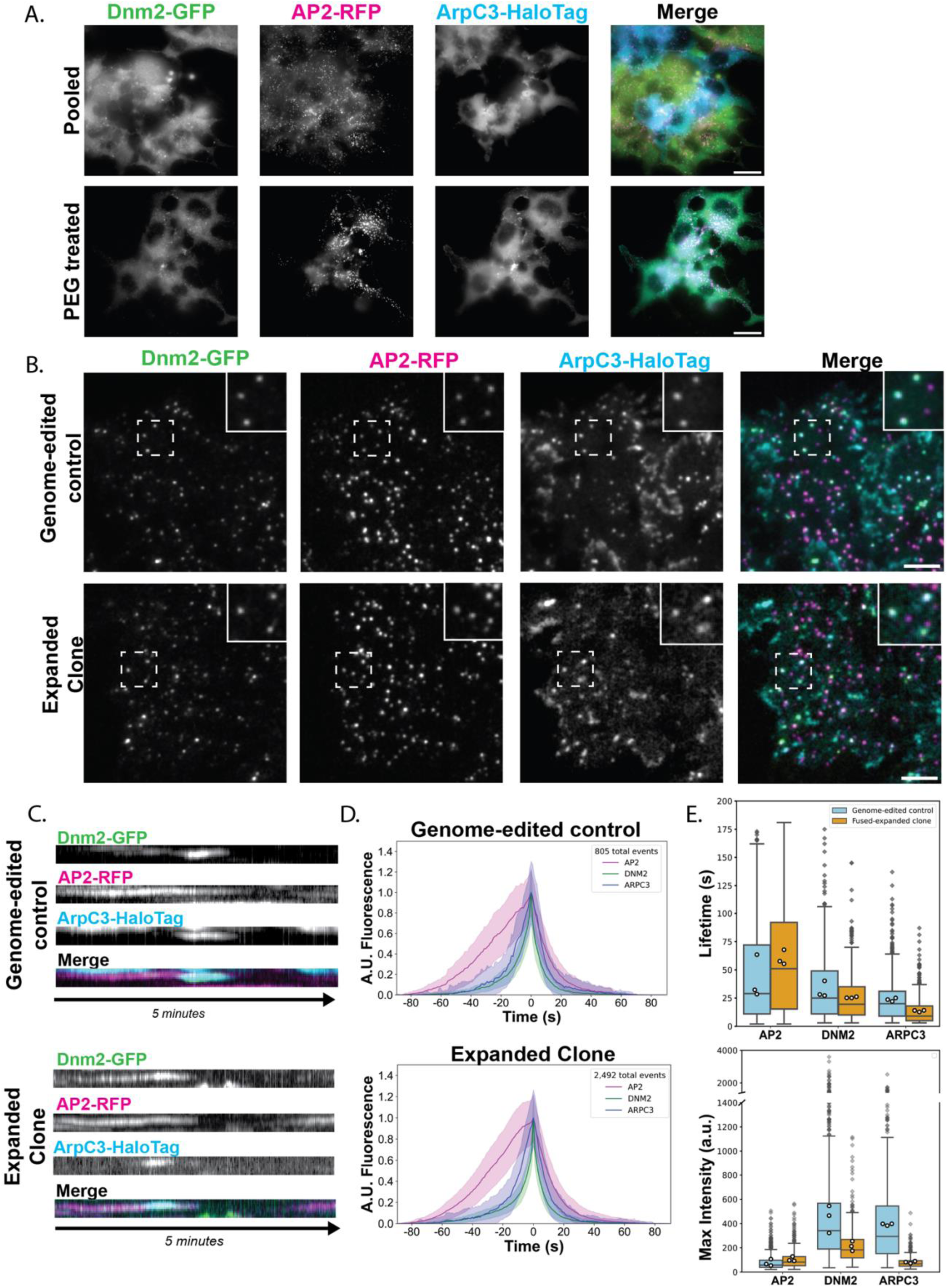
(A) Hi-Lo TIRF imaging of AP2-RFP; DNM2-GFP two-colored cell line co-cultured with AP2-RFP; ArpC3-HaloTag two-colored cell line (top). After treatment with PEG, cells express AP2-RFP, Dmn2-GFP, and ArpC3-HaloTag (bottom). Scale bar = 20 µm. (B) TIRF still image of CRISPR-Cas9 genome-edited cell line expressing fluorescent fusion proteins of three different colors (top), and a clonally expanded cell line generated from fusing two cell lines each expressing fusion proteins of two colors (bottom). Scale bar = 5 µm. (C) Kymographs showing the lifetime trace of a single CME event across a 5-minute movie. (D) Averaged intensity versus time plots of ArpC3-positive CME sites in genome-edited control cells and pooled fused clones. The lifetimes of the events plotted are between 20 and 180 seconds for 79.25% of the sites identified in the control and 88.72% of the sites identified in the fused clone. Events are aligned to the frames showing the maximum Dnm2 intensity (time = 0 sec). (E) Box and whisker plot comparing protein lifetimes for AP2, Dnm2, and ArpC3 in the genome-edited control cell line vs the fused, expanded iPSC clone cell line. The mean of each replicate is indicated by white circles. (F) Box and whisker plot comparing the fluorescent maximum intensity of AP2, Dnm2, and ArpC3 in the genome-edited control cell line and the fused, expanded iPSC cloned cell line. The mean of each replicate is indicated by white circles.

The protein localization at the basal lateral membrane of the fused cells was similar to that observed for the previously sequentially engineered hiPSC line that expresses AP2-RFP, Dnm2-GFP, and ArpC3-Halo (Figure 3B, (Jin et al., 2022). The ArpC3-Halo signal was also observed on the leading edge of the cell and in membrane ruffles, as a further indication that functional fluorescently tagged proteins are expressed in the fused cells. We next sought to clonally expand our fused three-color hiPSCs (Figure 2B). Through FACS we verified successful hybrid generation through observation of cells that were GFP positive and Halo positive.(Figure S2B). The fluorescence intensities of these hybrid cells matched those of the triple genome-edited lines, and these cells had increased DNA content when stained with Hoechst 3342 (Figure S2, C and D). Following clonal expansion of this fused population, cells contained only one nucleus but the number of chromosomes in each cell within the same clonal population was variable (Figure S2, E and F). The varying chromosome count is expected as it has been shown previously that hybrid cells at first have double the number of chromosomes, but they eventually lose chromosomes and approach a 2N state (Wollweber et al., 2000). Despite varying chromosome number, these fused clones maintained normal stem cell morphology and were able to survive multiple freeze-thaw cycles without undergoing spontaneous differentiation.

**Figure 3:**
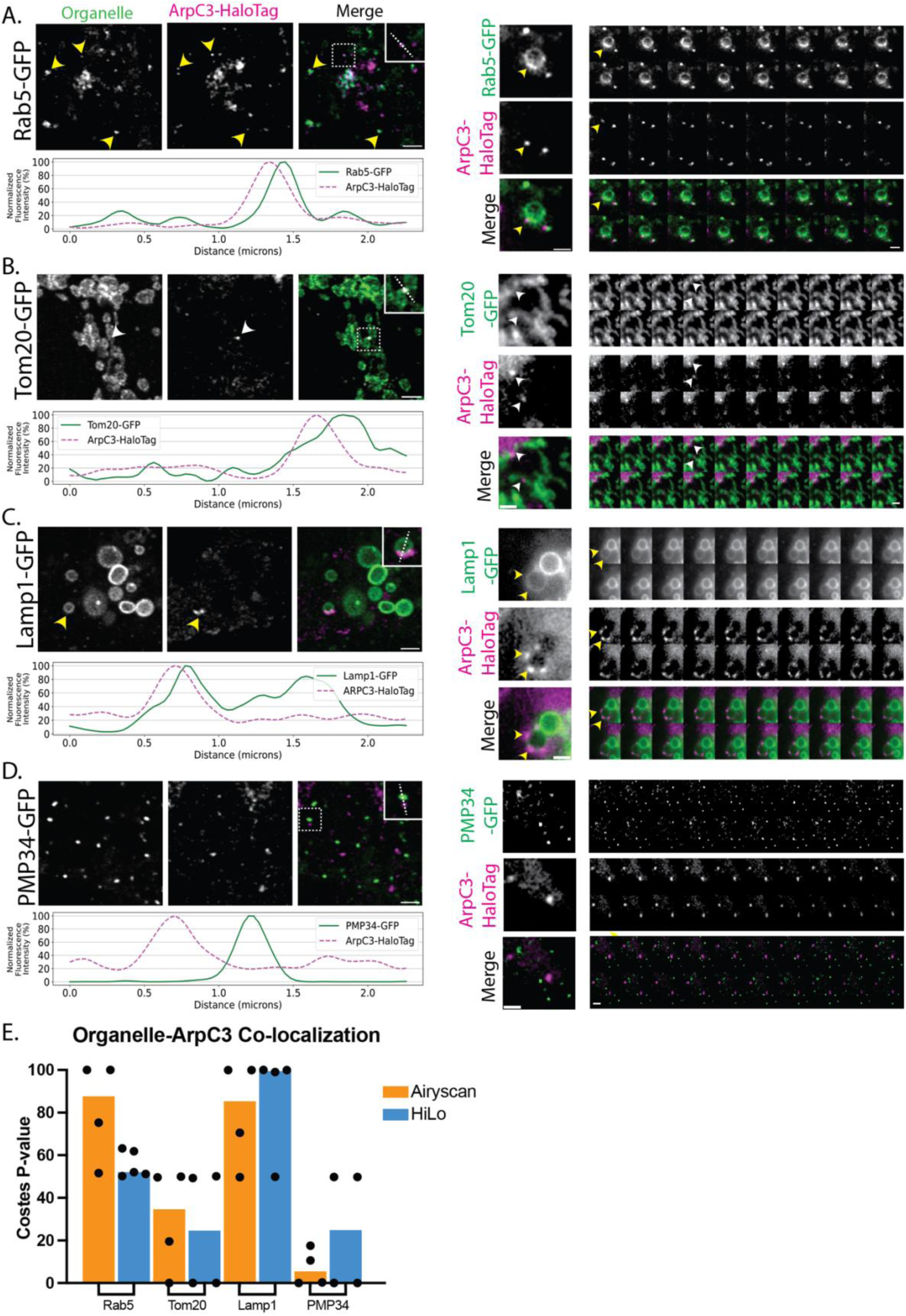
(A-D) Co-localization for ArpC3 and different organelle markers. Average intensity projection (z = 0.73 µm) from fixed samples. Yellow arrowheads indicate organelle co-localization with ArpC3 supported by Costes quantification. White arrowheads indicate observed co-localization that did not pass significant p-value using Costes quantification . Scale bar = 2 µm. Line scans (shown in the insets with dashed lines) were quantified and graphed (on the right). Merged colored montages from Hi-Lo TIRF movies are shown below, where each frame is 1 second for a total of 20 frames from a 2-minute movie. Scale bar = 2 µm. (E) Quantification of co-localization of ArpC3 with the different organelles for both fixed and live-cell imaging using Costes randomization.

We next sought to determine if endocytosis is normal in the clones of cells derived from the fused cells. In the expanded clones, the three tagged endocytic proteins all showed similar dynamics to those described previously in non-fused cells (Figure 2C, Movie 2). Using single particle tracking analysis for sites that had lifetimes between 20 and 180 seconds, we observed that protein recruitment and timing for each endocytic protein was similar to what was observed in the genome-edited control. AP2 arrives at the endocytic site first, followed by the arrival of the Arp2/3 complex and dynamin2 (Figure 2D). The non-fused, genome-edited control cells showed higher fluorescence intensity for ArpC3 and Dnm2 than the fused clones, which is expected because the fused clones presumably carry half of the number of edited Dnm2 alleles as the parent is homozygous for the edited allele, and one-quarter of the number of edited ArpC3 alleles as the parent is heterozygous for the edited allele (Figure 2E). Interestingly, if we multiply the maximum intensity of each protein by 4 and divide by the number of alleles tagged, we get more similar fluorescence intensity measurements for Dnm2 (control = ∼443.67 a.u., fused clone = 427.33 a.u.) and Arp2/3 (control = ∼392 a.u., fused clone = 322.67 a.u.). Based on these studies, we conclude that we succeeded in fusing and clonally expanding hiPSCs, and the resulting fused lines show similar endocytic protein dynamics to non-fused, genome-edited cells. These observations suggest that cell fusion can be useful for first-pass characterization of physiological interactions between genetically encoded fusion proteins.

### Systematic screen of endocytic protein-organelle interactions in fused hiPSCs

After establishing that endocytic proteins in fused hiPSCs exhibit normal dynamic behavior at the plasma membrane, we wondered if cell fusion could be used to identify novel functions for endocytic proteins. Clathrin-coated structures are known to associate with certain intracellular membranes, such as the surface of endosomes, where they are involved in recycling receptors back to the plasma membrane (Futter et al., 1998; Stoorvogel et al., 1996; Vassilopoulos & Montagnac, 2024). We wondered if other endocytic proteins have functions in various membrane-bound compartments within the cell. By fusing our hiPSC line expressing AP2-RFP and ArpC3-Halo to hiPSC lines edited to express GFP-tagged organelle markers (allencell.org/cell-catalog), Rab 5 (endosome), Tom20 (mitochondrion), Lamp1 (lysosome), PMP34 (peroxisome), Sec61 (ER), and sialytransferase 1 (Golgi), we screened for interactions between AP2 and Arp2/3 and the indicated organelles using fluorescence microscopy. Our results revealed both previously observed and novel co-localization between the query proteins and specific organelle markers (Table 1). We implemented Costes randomization to help us test the significance of any apparent colocalization, using 90% as the cut-off point for true co-localization (Bolte & Cordelières, 2006; Costes et al., 2004). Our results demonstrate that AP2 and ArpC3 most strongly co-localize with Rab5 and Lamp1, less so for Tom20, and little to no co-localization with PMP34 (Figure 3E & 4E). We first screened for co-localization between the query proteins and the early endosome marker Rab5 (Figure 3A & 4A, Movie 3). Early endosome morphology maintenance is known to depend on WASH, an Arp2/3 complex nucleating promoting factor (NPF), and the Arp2/3 complex (Duleh & Welch, 2010). WASH or Arp2/3 depletion through siRNA treatment result in elongated, tubulated early endosomes (Derivery et al., 2009). We observed that the Arp2/3 complex and Rab5 colocalize tightly and move with one another, and that endosome morphology is not perturbed in these fused cells (Figure 3A). Interestingly, we also observed AP2 at endosomes (Figure 4A). Other Adaptor Protein (AP) complexes, AP1, AP3, AP4, and AP5, but not AP2, have been shown to participate in intracellular trafficking (Sanger et al., 2019). We observe multiple instances of Rab5 co-localization with AP2 and hypothesize that this association might occur soon after an endocytic clathrin-coated vesicle is internalized from the plasma membrane as AP2 begins to disassemble from the vesicle, which transitions into an early endosome. Indeed, evidence from *in vitro* experiments suggests that AP2 uncoating from clathrin-coated vesicles (CCVs) is dependent on functional Rab5 (Semerdjieva et al., 2008). Our work allows us to visualize AP2 and Rab5-positive endosomes in live cells. Future experiments are required to analyze the dynamic and functional interplay between these proteins.

**Figure 4:**
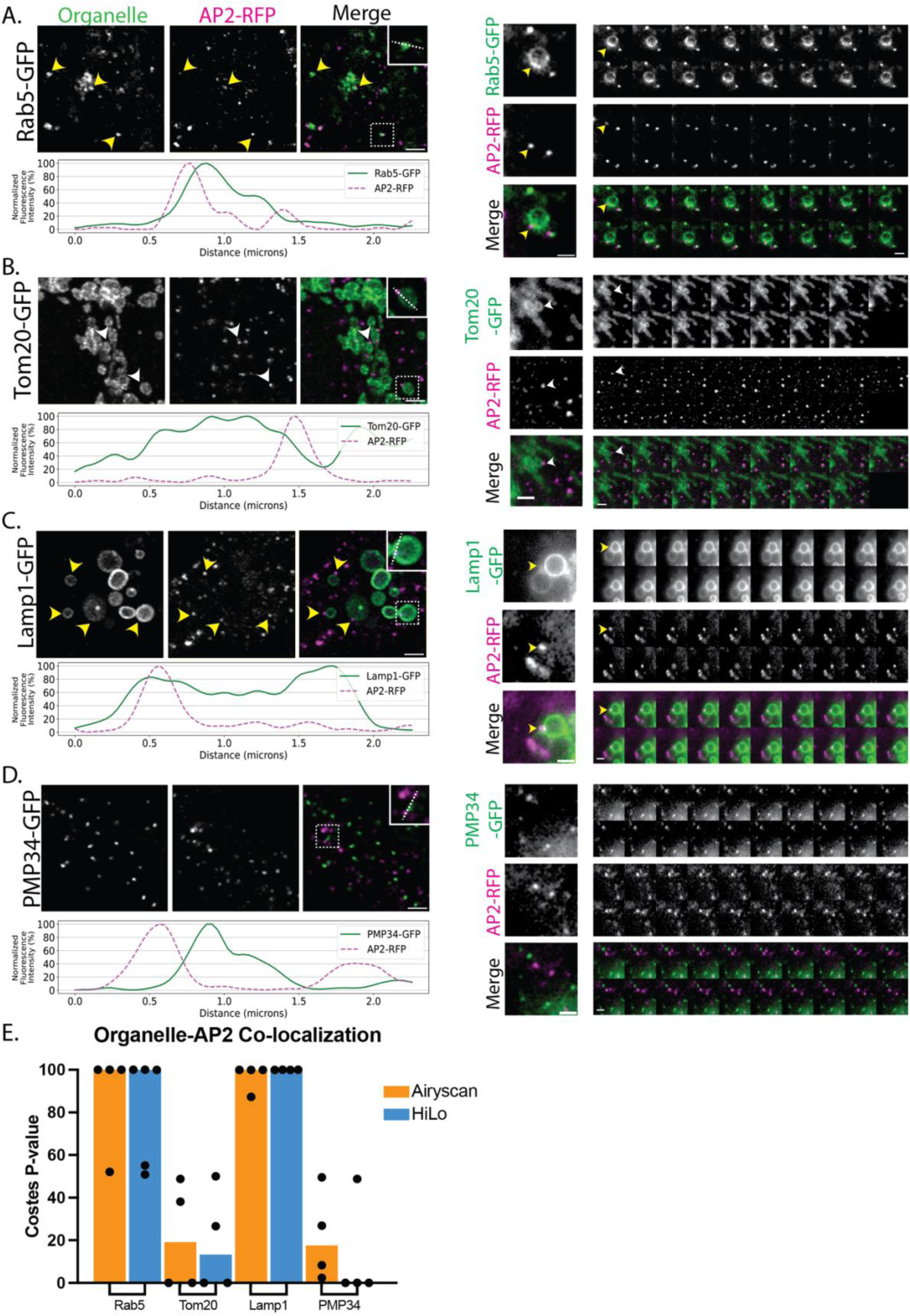
(A-D) Co-localization for AP2 and the different organelle markers. Average intensity projection (z = 0.73 µm) from fixed samples. Yellow arrowheads indicate organelle co-localization with AP2 supported by Costes quantification. White arrowheads indicate observed co-localization. Scale bar = 2 µm. Line scans (shown in the insets with dashed lines) were quantified and graphed (on the right). Merged colored montages from Hi-Lo TIRF movies are shown below, where each frame is 1 second for a total of 20 frames from a 2-minute movie. Scale bar = 2 µm. (E) Quantification of co-localization of AP2 with the different organelles for both fixed and live-cell imaging using Costes randomization.

**Table 1.** Results of cell fusion screen.

| Organelle | Marker | Co-localization with ArpC3 | Co-localization with AP2 |
| --- | --- | --- | --- |
| **Endosome** | Rab 5 | + | + |
| **Mitochondria** | Tom20 | + | + |
| **Lysosome** | Lamp1 | + | + |
| **Peroxisome** | PMP34 | - | - |
| **ER** | Sec61 | + | + |
| **Golgi** | Sialytransferase | - | - |

We next observed co-localization between the mitochondrial protein Tom20 and the Arp2/3 complex (Figure 3B, Movie 4). Previous research in HeLa cells reported that F-actin is recruited to the outer mitochondrial membrane to promote mitochondrial fission (Li et al., 2015). Consistent with this finding, F-actin has been reported to localize to mitochondria and treatments with compounds such as CK-666, an inhibitor of the Arp2/3 complex, led to complete dissociation of filamentous actin with mitochondria, suggesting that the Arp2/3 complex nucleates assembly of actin filaments on mitochondria (Moore et al., 2016). Our results confirm that the Arp2/3 complex associates with mitochondria. Line-scan measurements are consistent with the Arp2/3 complex associating with the mitochondrial membrane (Figure 3B). We also found that AP2 co - localizes with Tom20 (Figure 4B, right). Historically, CCVs have been proposed to either mature into early endosomes and follow the endolysosomal pathway or fuse with the trans-Golgi network (TGN) for sorting and packaging of proteins for intracellular transport (Cui et al., 2022). While the Costes randomization analysis did not indicate a strong correlation between AP2 and Tom20, we did observe a rare association between the two, which is a surprising finding, as mitochondria and the trans-Golgi network are separate compartments (Ford et al., 2021; Tábara et al., 2021). However, recent work challenged this conclusion, revealing that four-way contact sites link mitochondria, trans-Golgi vesicles, lysosomes, and ER membranes at sites of mitochondrial fission (Nagashima et al., 2020) . With a diverse mix of membranes found at these sites, it is possible to imagine that AP2 is also playing a role in establishing where mitochondrial fission sites arise.

We also fused the ArpC3/AP2 query cell line to cells expressing markers for the Golgi and the ER (Figure S3). We could not positively assess co-localization with the Golgi or the ER as the signal of the Golgi marker, ST6GAL1, and the ER marker, Sec61 was too heterogenous for us to confidently segment. Upon the completion of our screen, we decided to further analyze the relationship between AP2 and ArpC3 markers and lysosome markers.

### Association of actin and CME proteins with lysosomes

We further investigated the interactions between CME proteins, actin, and lysosomes. First, the identity of the LAMP1-marked compartments that AP2 and the Arp2/3 complex associated with was confirmed by staining hiPSC expressing Lamp1-GFP with antibodies against several lysosome-associated proteins and consistently observing robust colocalization (Fig S.4A). Next, we utilized one of the tested antibodies, which recognized TMEM192, to verify the association between AP2 and lysosomes in a hiPSC line expressing AP2-TagRFP-T and DNM2-tagGFP2 that had not undergone cell fusion (Figure 5A top row). We additionally stained these cells with antibodies against clathrin heavy-chain and observed colocalization of all three clathrin-mediated endocytosis proteins in cortical foci around the surface of the lysosome (Figure 5A, top row). Since ArpC3 was also observed in association with the cortex of lysosomes, we also stained the same hiPSC line with fluorescent phalloidin, which has high affinity for filamentous actin. The phalloidin stain revealed cortical actin puncta that co-localize with CME proteins on the lysosomal surface (Figure 5A, bottom row). Collectively, we observed three major endocytic proteins, AP2, DNM2 and clathrin, as well as actin, localized in punctate structures on the cortex of lysosomes in unfused iPSCs.

**Figure 5:**
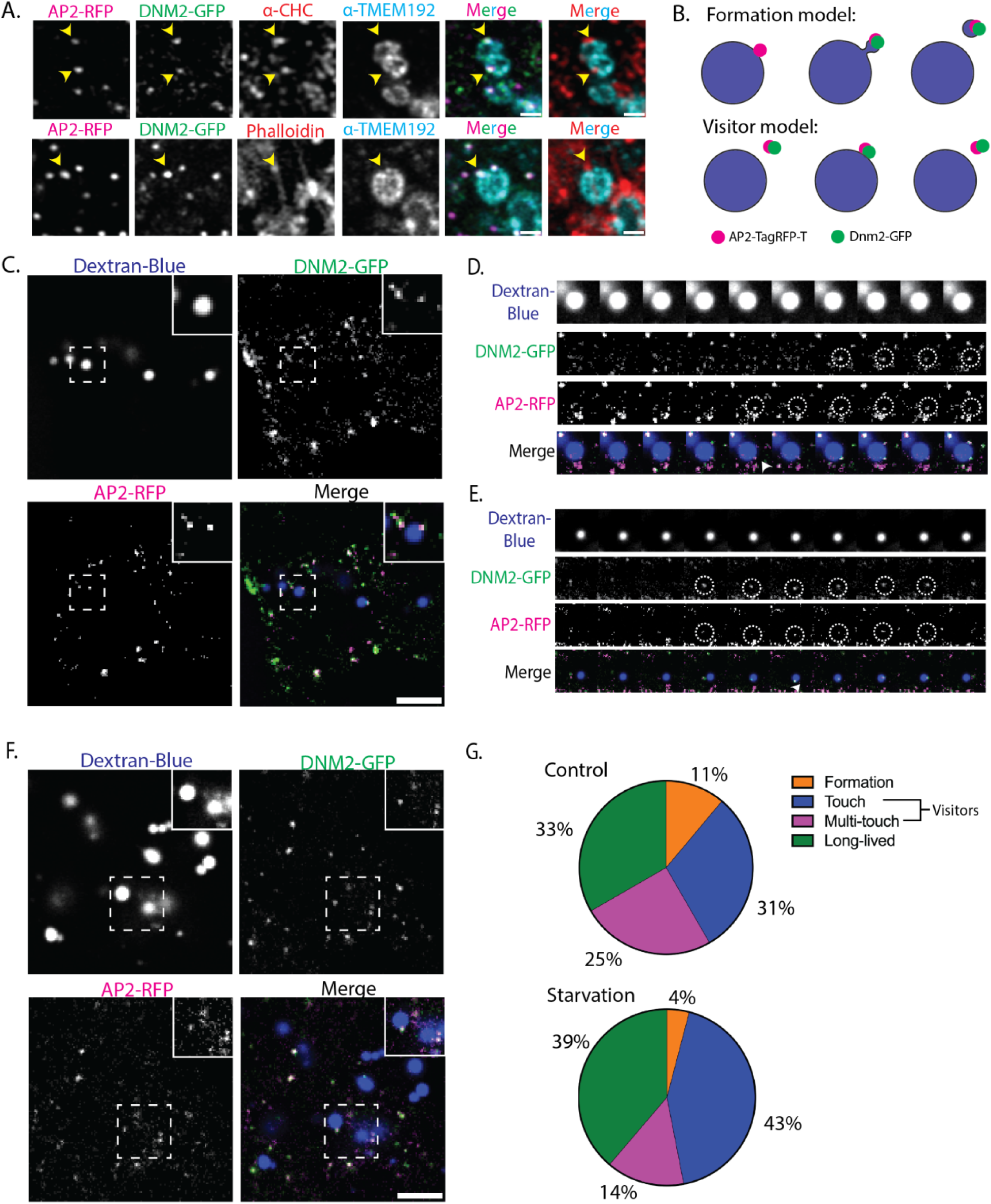
Image analysis of interactions between AP2, actin, and endolysosomal organelles. (A) Single z-slice (146 nm slice thickness) of fixed genome-edited hiPSCs expressing AP2-RFP and Dnm2-GFP stained with anti-clathrin Heavy Chain and anti-TMEM192 lysosomal protein (top row) to demonstrate co-localization of CME proteins with the lysosome (yellow-arrowheads), and phalloidin and TMEM192 (bottom row) to demonstrate co-localization of CME proteins, actin, and the lysosome (yellow-arrowheads). Scale bar = 2 µm. (B) Schematic modeling of two possible types of events involving CME proteins on the lysosome surface. (C) Single z-slice (0.4 µm) from live-cell image of genome-edited hiPSCs expressing AP2-RFP and Dnm2-GFP and incubated with 10 kDa Dextran-Cascade Blue. The inset shows a single lysosome with both AP2 and DNM2 puncta. Scale bar = 5 µm. (D) Montage of CME-like event on the lysosome surface imaged at 1 frame/3 seconds. (E) Montage of a likely pre-formed “visitor” arriving at the lysosome surface. (F) Single z-slice (0.4 µm) from live cell image of hiPSCs after 8 hours of starvation. The inset shows multiple lysosomes with AP2 and Dnm2 puncta. Scale bar = 5 µm (G) Quantitative comparison of control vs starved hiPSCs, counting how many of each type of event was observed. Thirty-six events were observed under control conditions, and 49 events were observed under starved conditions.

Confirming the presence of actin and several CME proteins on the surface of lysosomes, we wondered whether a CME-like process might be occurring on the lysosome surface. To test this possibility, we posited two alternative models of lysosome-associated endocytic protein dynamics, the “formation” and “visitor” models. The formation model posits that endocytic protein assembly is nucleated on the surface of the lysosome, perhaps culminating in a budding event to release a vesicle (Figure 5B, top). The visitor model posits that the endocytic proteins arrive as preformed structures, such as an endocytic vesicle that contacts the lysosomal surface (Figure 5B, bottom). To distinguish between these two possibilities, we utilized live-cell imaging to determine whether endocytic proteins gradually accumulated on the lysosomal surface, consistent with the formation model, or appeared suddenly, consistent with visitor model. To this end, hiPSCs expressing AP2-RFP and DNM2-GFP were incubated with Cascade-blue conjugated Dextran, a fluorescent molecule that accumulates in lysosomes, and imaged using a Nikon NSPARC microscope (Figure 5C and S.4B and C). A variety of events were observed by taking high-speed multi-channel z-stack movies in which 3-channel 3.6 µm z-stacks were acquired at 0.3 frames/second. We observed that about 11% (total n = 36) of the associations involved the gradual accumulation of endocytic proteins, consistent with the formation model. Moreover, we observed the ordered recruitment of CME proteins in which AP2 arrives first, followed by a burst of DNM2, reminiscent of CME events (Figure 5D, Movie 6). In the majority of events (56%), AP2 and DNM2 arrived at the lysosomal surface concurrently, as preformed foci that contacted the lysosome surface once or more than once, consistent with the visitor model (Figure 5E and G, Movie 7). Lastly, about 33% of the associations lasted for the full duration of the 2-minute movie acquisition. We categorized these sites as long-lived events, similar to the static CME sites that often are observed. Our findings suggest that the “visitor” model describes many of the observed CME protein associations with lysosomes.

Several studies reported the association of CME proteins with autolysosomes, which recycle lysosomal components during autophagy. These publications collectively support a model in which a CME-like process plays a role in lysosome regeneration after starvation-induced autophagy in a pathway called autophagic lysosome reformation (ALR) (Dai et al., 2019; Du et al., 2016; Rong et al., 2012; Schulze et al., 2013). We therefore determined whether endocytic protein dynamics on the lysosomal surface would shift under starvation conditions when autophagy is upregulated. To determine if CME protein dynamics associated with lysosomes change in response to autophagy induction, we performed an 8-hour starvation of the hiPSCs expressing AP2-tagRFP-T and DNM2-GFP (Figure 5F, Movie 8). We qualitatively categorized the events as formation or visitor events (Figure 5G). The majority (57%) of observed events fit the visitor model, with CME proteins arriving at the lysosome as pre-formed puncta. However, ∼4% of the events showed a gradual accumulation of endocytic proteins reminiscent of a canonical CME site formation at the plasma membrane. Thus, we conclude most endocytic protein puncta associated with lysosomes show visitor-like dynamics and likely do not represent CME-like budding events, regardless of whether the cells had been subjected to starvation or not. It is possible that the visitor events we observed are the trafficking of Lamps to the lysosome from the plasma membrane (Janvier & Bonifacino, 2005). Further analysis is required to distinguish between these two possibilities.

## Discussion

For decades, cell fusion has served as a powerful method for exploring biological mechanisms. Rao and Johnson fused HeLa cells at different stages of the cell cycle to gain insights into the regulation of DNA replication and mitosis (Rao & Johnson, 1970). Harris and Klein fused normal cells with cancer cells to investigate oncogene dominance, leading to the discovery of tumor suppressor genes (Harris et al., 1969; Lipsick, 2020). Gokhman et. al. fused chimpanzee and human induced pluripotent stem cells to probe cis-regulatory element function in gene regulation (Gokhman et al., 2021). Here, we used cell fusion among genome-edited human cell lines to rapidly screen for novel protein functions. Libraries of genome-edited cell lines are becoming valuable resources. However, while it can be desirable to survey combinations of alleles generated in these lines as is routinely done for model organisms like yeast and fruit flies that can be mated to each other, creating new genomic combinations quickly with minimal impact on expression levels has not been possible.

In this study, we utilized cell fusion to generate clonally expandable, genome-edited hybrid cell lines that express new combinations of proteins tagged with fluorescent proteins. This approach leveraged a collection of isogenic, genome-edited hiPSC lines, each expressing from its endogenous chromosomal locus, an organelle protein marker tagged with GFP, significantly reducing the time needed to explore spatio-dynamic relationships between proteins and organelles.

To determine how cell fusion impacts cellular function, we used CME pathway dynamics as a functional readout, as defects in endocytic protein expression result in irregular endocytic protein spatio-dynamics. We found that cell fusion preserved normal protein localization and normal dynamics of the CME pathway. We did, however, observe genetic instability in isolated clones of fused cells, as indicated by karyotyping. Thus, while cell fusion is a powerful approach for a preliminary screen to identify previously unobserved functional relationships, it is important to verify and further investigate those inferred functional relationships in unfused cells.

To this end, we further investigated the nature of the observed interaction between CME proteins and lysosomes. Following our cell fusion screen, we observed co-localization between CME proteins and LAMP1 in unfused, healthy hiPSCs. We observed CME protein dynamics on the lysosome surface using high-speed volumetric imaging, some reminiscent of CME-like budding events. However, most were categorized as visitor events by CME vesicles in normal and starvation conditions. Previous work has observed co-localization of CME proteins, including AP2, clathrin, dynamin, and actin with autolysosomes during starvation-mediated autophagy. Still, this study is the first to demonstrate co-localization of multiple endocytic proteins on the lysosome surface and measure endogenous endocytic protein dynamics at the lysosome in healthy cells. These observations may demonstrate that these rare events are part of the ALR pathway during non-starvation-mediated autophagy in hiPSCs, or conversely, another part of the endolysosomal pathway. The use of cell fusion allowed us to quickly screen for co-localization of multiple fluorescently tagged proteins not previously co-expressed without the use of transient transfection or multiple rounds of gene editing. The rapid nature of this approach enabled us to follow up on an observed interaction and plan experiments more intuitively using existing tools in the lab. We believe that cell fusion serves as a suitable tactic for investigating novel protein localization and dynamics.

## Supporting information

Thresholding Information

## Acknowledgments

We thank Alison Killilea, the Stem Cell Facility Director at UC Berkeley, for providing the reagents and cell lines for this project. We thank the Allen Institute for Cell Science for the hiPSCs used in the cell fusion screen. We thank Meiyan Jin, Yidi Sun, Jonathan Wong, and our anonymous reviewers for their critical reading of the manuscript. This research was conducted with US Government support, under NIH Grant R35GM118149 to D.G.D.

## Author Contributions

S.L.S, Y.I, and A.M. performed research, analyzed data; S.L.S, Y.I, A.M., and D.G.D designed the research and wrote the manuscript.

## Methods & Materials

### Cell Culture

The WTC10 hiPSC line was obtained from the Bruce Conklin lab at UCSF. The hiPSCs were cultured on Matrigel (hESC-Qualified Matrix, Corning #354277) coated dishes in StemFlex medium (Thermo Fisher #A3349401) with Penicillin/Streptomycin in 37°C with 5% CO_2_. Cultures were passaged with Versene (Thermo Fisher #15040066) twice a week. SK-MEL-2 cells were cultured in DMEM F12 + GlutaMAX (Thermo Fisher #10565018) with 10% FBS and Penicillin/Streptomycin in 37°C with 5% CO_2_. Cultures were passaged with 0.25% Trypsin-EDTA (Thermo Fisher #25200114) twice a week.

### Genome-editing

The ARPC3 gene was edited in WTC10 hiPSCs in an AP2M1 dual tagged TagRFP-T background, which was previously edited. The Cas9-sgRNA complex electroporation method was used to edit the ARPC3 gene to insert the HaloTag. *S. pyogenes* NLS-Cas9 was purified in the University of California, Berkeley QB3 MacroLab. The ARPC3 sgRNA (CCTGGACAGTGAAGGGAGCC) was purchased from IDT. Gibson assembly (New England Biolabs) was used to construct the donor plasmid containing ARPC3 5’homology-ggatccggtaccagcgatccaccggtcgccacc-HaloTag-ARPC3 3’ homology. One week after electroporation (Lonza, Cat #: VOH-5012) of the Cas9-sgRNA complex and donor plasmid, the HaloTag positive cells were single cell sorted using a BD FACSAria Fusion Flow Cytometer (BD Biosciences) into Matrigel-coated 96-well plates with StemFlex medium with ROCK Inhibitor. Clones were confirmed by PCR and Sanger sequencing of the genome DNA locus around the insertion site. One allele of ARPC3 was tagged with HaloTag in the hiPSC lines used.

### Cell Fusion

The protocol for cell fusion was adapted from Agoglia et al., 2021. PEG 1500 (Sigma-Aldrich #10783641001) was used to mediate cell fusion of either SK-MEL-2 cells or hiPSCs. Briefly, each parent cell line was co-cultured until well reached 80% confluence in a 6-well plate. Media was aspirated, washed one time with corresponding media (either SK-MEL-2 or hiPSC specific media), and 1 mL of PEG 1500 was pipetted to each well and incubated at 37°C for 2 minutes. The cells were washed twice gently with 1 mL of corresponding media. A more detailed protocol for each cell type is provided below.

### Clonal Expansion

The two SK-MEL-2 parental genotypes (Clta-TagRFP-T and DNM2-eGFP) were washed once with 1X Dulbecco’s Phosphate Buffered Saline (DPBS) and then treated with Trypsin for 3 minutes at 37°C. Cells were pelleted and resuspended in SK-MEL-2 media. The cells were plated at equal concentration (∼0.5 million cells each) and co-cultured together in a 6-well plate. Media was changed every other day until 70-80% confluent.

The two hiPSC parental genotypes (AP2-TagRFP-T; ArpC3-HaloTag and AP2-TagRFP-T; Dnm2-tagGFP2) were washed once with 1XDPBS and then treated with Accutase (Thermo Fisher #A1110501) for 3 minutes at 37°C. The cells were resuspended in StemFlex, pelleted and resuspended in StemFlex + 10 µM ROCK Inhibitor. The two cell lines were plated in a Matrigel coated 6-well plate containing StemFlex with ROCK Inhibitor at equal concentrations of ∼1×10^6^ cells per genotype. Media was changed to StemFlex without ROCK Inhibitor the following day.

Once the cells reached ∼80% confluency, the cells were washed with media and then incubated with 1 mL of PEG 1500 for 2 minutes at 37°C. After, the fusogen was removed and the cells were gently washed twice with corresponding media (either SK-MEL-2 or hiPSC media) and then kept in media for 24 hours. For hiPSCs this media was StemFlex with ROCK inhibitor for 24 hours. The unfused, pooled cells were used as a control. After 24 hours, the media was replaced with normal media without ROCK inhibitor .

The fused cells recovered for two days. Prior to FACS, the hiPSCs were incubated in StemFlex medium with 100 mM JF635-HaloTag ligand (Grimm et al., 2017) for 1 hour. The unbound ligand was washed away with two 10-minute washes. For FACS preparation, the cells were washed once with DPBS and then treated with Accutase or Trypsin-EDTA for 3 minutes at 37°C. The cells were spun down and resuspended in medium. The cells were sorted for positive signal in the Allophycocyanin (APC), Fluorescein isothiocyanate (FITC), and Phycoerythrin (PE) channels using a BD FACSAria Fusion Flow Cytometer (BD Biosciences) into either 96-well plate containing DMEM medium or Matrigel-coated 96-well plate containing StemFlex medium with ROCK inhibitor. The cell media was changed to cell-type specific media after three days, and subsequently changed every other day. Clones were confirmed by PCR and Sanger sequencing of the genome DNA locus around the insertion site, as well as western blot analysis. Karyotyping was performed on the hiPSC parental cell lines and clonally expanded fused cells were performed by KaryoLogic.Inc.

#### Overexpression via Transient Transfection

250uL of OptiMEM I Reduced Serum (Thermo Fisher #31985070) media warmed to 37°C was added to a 1.5mL microcentrifuge tube. 2ug of the plasmid containing the protein for overexpression was added and mixed. 7.5 uL of Mirus Bio TransIT-LT1 (MIR 2304) reagent was added to the tube and incubated for 30 minutes. TransIT-LT1:DNA complex was pipetted drop wise to a 70-80% confluent well of SK-MEL-2 cells in a 6 well plate, plate was rocked for even distribution. Plate was incubated for 24-72 hours and then cells were harvested for flow cytometry or imaging.

### Organelle Screen Fusion

The two hiPSC parental genotypes (AP2-TagRFP-T;ArpC3-HaloTag and Allen Institute Cell line with a GFP organelle marker) were washed once with DPBS and then treated with Accutase for 3 minutes at 37°C. The cells were pelleted and resuspended in StemFlex with ROCK Inhibitor. The two cell lines were seeded in a Matrigel coated 4-well chambered cover glass (Cellvis) containing StemFlex with ROCK Inhibitor at equal concentrations of ∼5×10^4^ cells per genotype. After 24 hours, the cells were washed with media and then incubated with PEG 1500 for 2 minutes at 37°C. After, the fusogen was removed and the cells were gently washed twice with StemFlex and then kept in StemFlex medium with ROCK inhibitor for 24 hours. Unfused, pooled cells were used as a control. The media was exchanged with StemFlex medium after 24 hours, and the cells were prepared for imaging the following day.

### Dextran treatment

Lyophilized Dextran Cascade Blue 10 kDa (Thermo Fisher #D1976) was resuspended in 1 mL of ddH_2_O to 25 mg/ml. Aliquots of dextran were diluted in StemFlex media and kept at -20°C at a concentration of 8 mg/mL. Cells were incubated with dextran (4 mg/mL) for 48 hours at 37°C. The day of imaging, cells were briefly washed with StemFlex 3 times before the addition of imaging media (DMEM/F12 + 10X StemFlex Supplement + 10 mM HEPES buffer).

### TIRF Live-Cell Imaging

Two days before imaging, the hiPSCs were seeded onto Matrigel-coated cover glasses in 4-well chambers (Cellvis). Two hours before imaging, the cells were incubated with 100 mM JF635-HaloTag ligand in StemFlex medium for 1 hour. After incubation, the cells were washed with StemFlex medium twice for 10 minutes at 37°C. SK-MEL-2s were instead seeded onto cover glasses in 4-well chambers with DMEM medium without phenol red and imaged the next day. The cells were imaged using a Nikon Ti2-E inverted microscope. During imaging, the cells were maintained at 37°C with a stage top incubator (Oko Labs) in DMEM/F12 + 10X StemFlex Supplement + 10 mM HEPES buffer. Images were acquired using a Nikon 60X CFI Apo TIRF objective (1.49 NA) and an Orca Fusion Genn III sCMOS camera (Hamamatsu) at 1x magnification using the Nikon NIS Elements software. Using a LUNF 4-line laser launch (Nikon Instruments) and iLas2 TIRF/FRAP module (Gataca Systems) total internal reflection fluorescence (TIRF) illuminated a single focal plane of the field and was imaged every second for 5 minutes. Hi-Lo TIRF illuminates a single focal plane with a more perpendicular TIRF angle to capture intracellular signal.

### Airyscan Fixed-Cell Immunofluorescence (IF) Imaging

Two days before imaging, the hiPSCs were seeded onto Matrigel-coated cover glasses in 4-well chambers (Cellvis). Two hours prior to imaging, the cells were incubated with 100 mM JF635 - HaloTag ligand in StemFlex medium for 1 hour and 5 µg/ml Hoechst 33342 for 30 minutes. After incubation, the cells were washed with StemFlex medium twice for 10 minutes at 37°C. The cells were washed once with 1 X PBS and then fixed with ice-cold fixation buffer (4% Paraformaldehyde in 30 mM MES pH6.1, 450 mM NaCl, 15 mM EGTA, 15 mM Glucose, 15 mM MgCl_2_ (CS Buffer) for 20 minutes at room temperature. Following fixation, cells were washed 3 times for 5 minutes with room temperature PBS pH 7.4. After washing, the cells were incubated in blocking buffer [3% (w/v) BSA and 0.1%(w/v) Saponin in PBS pH 7.4] for 40 minutes at room temperature. The blocking buffer was removed, and the cells were incubated in primary antibodies overnight at 4°C. The following day cells were washed 3 times for 10 minutes each in a 10:1 dilution of PBS and blocking buffer at room temperature. The secondary antibody was diluted into blocking buffer and incubated at room temperature for 2 hours. After incubation, the cells were washed twice with 10:1 PBS:Blocking buffer, followed by 3 10-minute washes in PBS. The cells were kept at 4°C until ready to image.

### Starvation Assay

Cells were first plated in an imaging dish and left to recover for 24 hours. The cells were treated with Dextran in StemFlex for 40 hours, then the media was substituted with Dextran in Hanks Balanced Salt Solution (Thermo Fisher #14025-092) for 8 hours. The cells were washed quickly three times with Hanks Balanced Solution without Dextran, then placed in Hanks Balanced Solution + 10 mM HEPES buffer. Cells were imaged immediately after.

### NSPARC

The cells were imaged on a Nikon Ti2-E inverted microscope with a Prior Queensgate stagetop Piezo Z drive. The cells were maintained at 37°C with a stage top incubator (Oko Labs) in DMEM/F12 + 10X StemFlex Supplement + 10 mM HEPES buffer. Images were acquired using a Nikon Apo TIRF 60x Oil DIC N2 (1.49 NA) and AXR laser-scanning confocal with resonant scanner and an NSPARC ISM (Image Scanning Microscopy) detector at 1x magnification using the Nikon NIS Elements software. Using a LUA-S4 4-line laser launch (Nikon Instruments) and imaged z-stacks with 0.4 µm step size at 0.33 frames/second.

### Organelle Segmentation/Costes Randomization

In order to perform Costes Randomization using Fiji/ImageJ, masks were generated for the images collected using Airyscan and HiLo microscopy. All of the specific parameters used for each organelle and imaging modality can be found in the Supplemental Information. Briefly, first a rolling ball background subtraction was applied, followed by a Gaussian Blur filter. Then, a threshold image was generated using the automated algorithm feature in Fiji/ImageJ. Each threshold algorithm was manually selected based on how successfully the mask matched the organelle signal for each organelle before moving forward with batch processing. The images were then cropped to generate a smaller region of the cell to reduce the noise from neighboring non-fused cells. The JACoP co-localization plugin was used to determine the Coste’s randomization p-value of the cropped, masked images. For the Airyscan images, a 16-pixel block was used and for the HiLo images an 8-pixel block was used.

## Code Availability & Image Analysis tools

https://github.com/DrubinBarnes

### Antibodies

| Antibody | Catalog Number | Concentration Used | Experiment |
| --- | --- | --- | --- |
| anti-mouse 647 | Invitrogen (#A-31571) | 1:400 | IF |
| anti-rabbit 647 | Invitrogen (#A-21443) | 1:400 | IF |
| anti-Clathrin heavy chain (X22) | Invitrogen (#MA1-065) | 1:250 | IF |
| anti-LAMP1 | Cell Signaling Technologies (#15665) | 1:100 | IF |
| anti-TMEM-192 | Abcam (#ab185545) | 1:100 | IF |
| anti-LAMTOR4 | Cell Signaling Technologies (#12284) | 1:300 | IF |
| anti-RAGA | Cell Signaling Technologies (#4357) | 1:300 | IF |
| anti-mTOR | Cell Signaling Technologies (#2983) | 1:400 | IF |
| anti-RAGC | Cell Signaling Technologies (#3360) | 1:300 | IF |
| CellMask Deep Red | Invitrogen (#C10046) | 1:1,000 | Live cell imaging |

## Supplemental Figures

**Figure S1:**
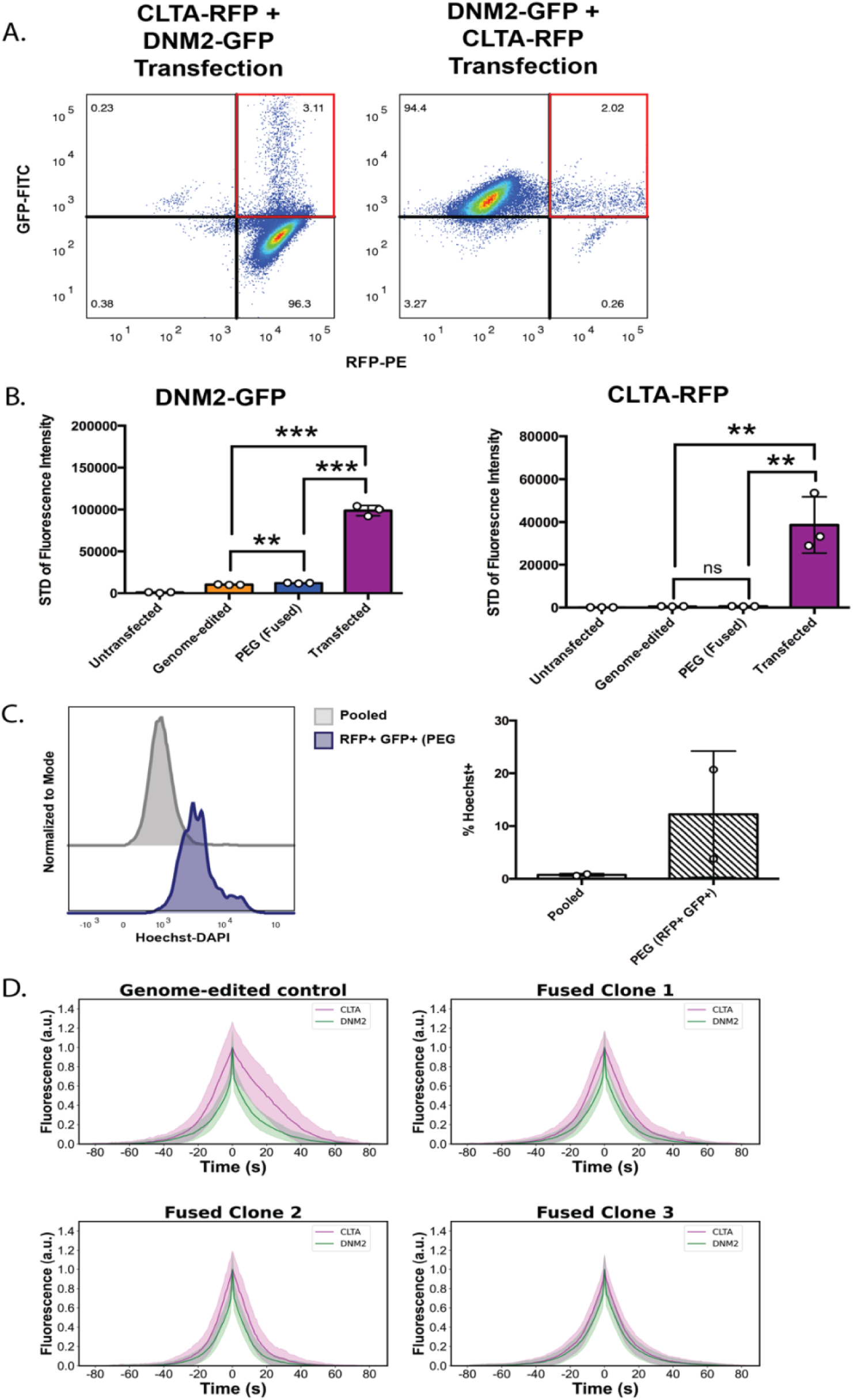
(A) Flow cytometry data from the cell fusion experiment discussed in Figure 1D. Signal from the PE laser (RFP) is on the x-axis, and signal from the FITC laser (GFP) is on the y-axis. The first plot shows data for the genome-edited Clta-RFP parent cell line overexpressing DNM2-GFP, and the second plot shows data for the genome-edited DNM2-GFP parent cell line overexpressing Clta-RFP. Quadrants are gated to measure the enrichment of a new GFP+ or RFP+ population highlighted in the red boxes in the plot as a result of overexpression. (B) Quantification of the standard deviation from Figure1D of fluorescence intensity for both GFP-FITC and RFP-PE across 3 replicates is shown in the bar graphs. (C) Histogram of Hoechst-DAPI signal for pooled Clta-RFP and DNM2-GFP parental cell lines and the RFP+ GFP+ population from the fused condition. Quantification of the Hoechst+ population (cells with Hoechst signal beyond the peak of the pooled cells) in the pooled and the RFP+ GFP+ fused population conditions across 3 replicates is shown in the bar graph. (D) Cohort plots of successively genome-edited, dual-colored cells, three examples of clonally expanded dual-colored cells after fusion to show the intensity of clathrin (magenta) and dynamin2 (green) over time during CME events in 3 replicates.

**Figure S2:**
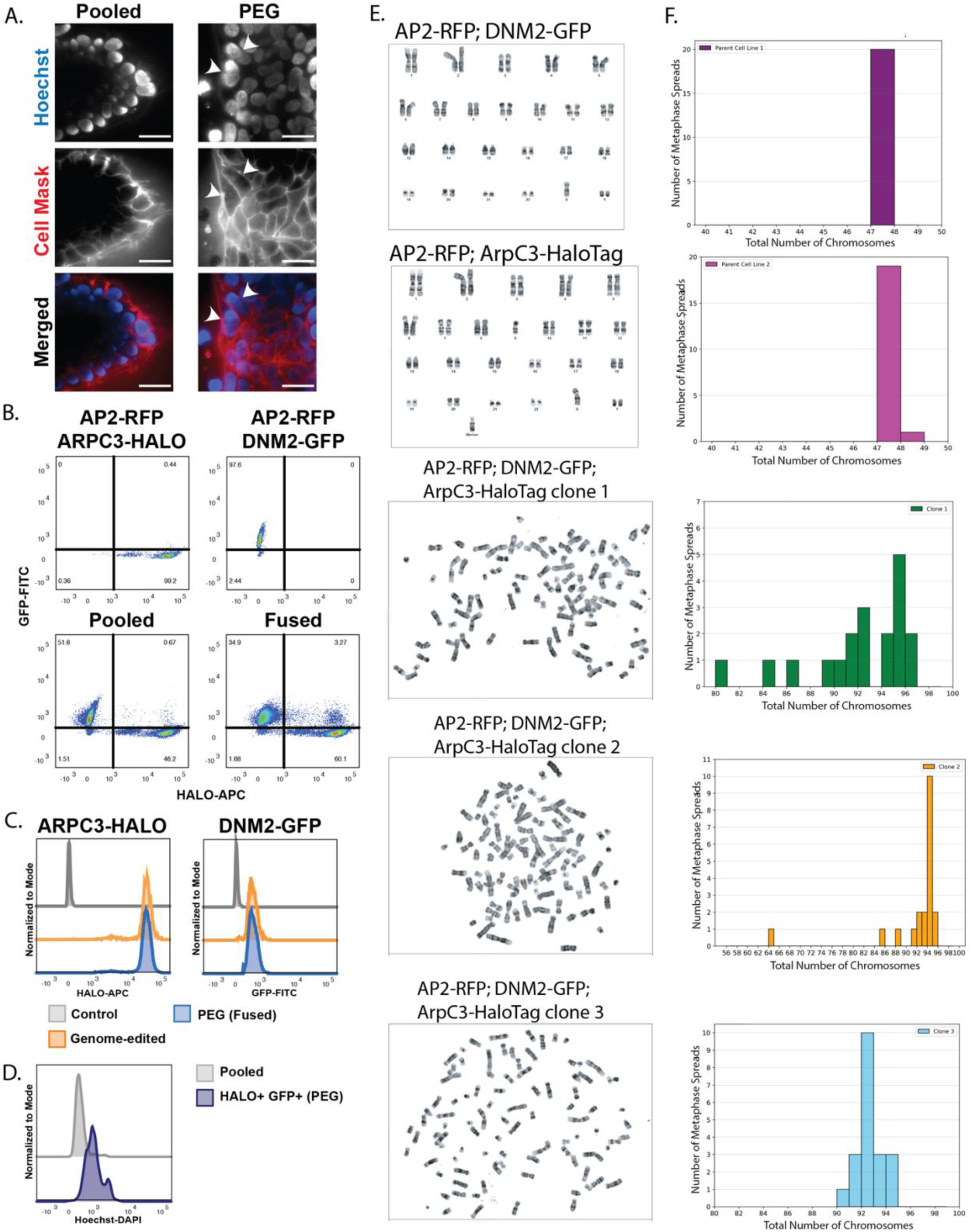
(A) TIRF still images on hiPSCs showing AP2-RFP; DNM2-GFP dual colored cell line co-cultured with AP2-RFP; ArpC3-HaloTag dual colored cell line without fusogen where the nuclei (blue) are clearly separated by the cell membrane (red), and the cell lines co-cultured with fusogen where some cells have multiple nuclei within one cell membrane marked by white arrowheads. Scale bar = 30 µm. (B) Flow cytometry data from the sorting of fused hiPSCs. The signal from the APC laser (HALO) is on the x-axis, and the signal from the FITC laser (GFP) is on the y-axis. The first plot shows data from the AP2-RFP; ArpC3-HaloTag parent cell line, the second plot shows data from the AP2-RFP; Dnm2-GFP parent cell line, the third plot shows data from the two parent cells co-cultured without fusogen, and the fourth plot shows data from the two parent cells co-cultured with fusogen. Quadrants are gated to measure the enrichment of a new APC+ GFP+ population in the plot, indicating co-culture and fusogen, and were used as a gate for sorting. (C) Histograms representing ArpC3-Halo and Dnm2-GFP signal for cells lacking HALO/GFP, genome-edited cells with the HALO/GFP to have a positive signal, and PEG treated HALO+ GFP+ cells. (D) Histogram showing data for the Hoechst-DAPI signal for pooled AP2-RFP; ArpC3-HaloTag and AP2-RFP; Dnm2-GFP parental cell lines and the HALO+ GFP+ population from the fused condition. Hoechst+ gating was done as in Figure S1 and used for sorting. (E) Cytogenetic analysis was performed on twenty G-banded metaphase spreads by Karyologic Inc. It appears that both parents carry an extra Y chromosome, and the AP2-RFP; ArpC3-HaloTag cell line lost a copy of chromosome 9 (top row). These cells nevertheless demonstrated normal CME dynamics, did not spontaneously differentiate, and underwent multiple freeze/thaw cycles throughout the course of this study without noticeable adverse effects on survival or physiology. The remaining karyotypes are from three different isolated single clones after two-rounds of FACS sorting. The first round of FACS sorting generated one successful clone that survived expansion, so we completed a second round of FACS sorting to isolate more single-cell clones for expansion. (E) Corresponding histogram of chromosome count from the 20 metaphase spreads. Each bar has a bin size of one.

**Figure S3:**
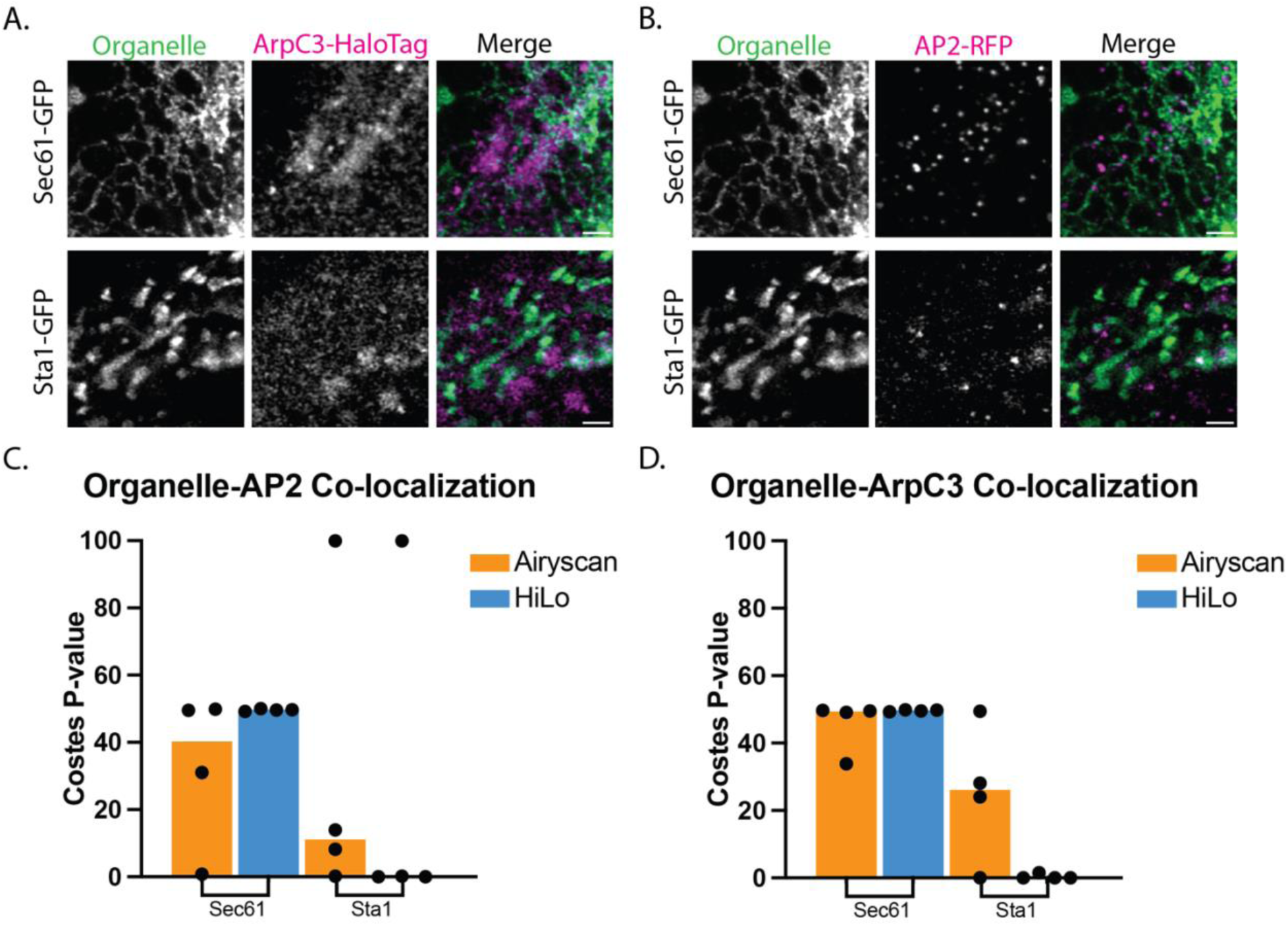
(A and B) Average intensity projection (z = 0.73 µm) from fixed samples of AP2-RFP; ArpC3-HaloTag cell line fused with a cell line expressing Sec61-GFP endoplasmic reticulum marker (top row) or Sta1-GFP (bottom row). Scale bar = 2 µm. (C & D) Quantification of co-localization of AP2 and ArpC3 with the Sec61 and Sta1 for both fixed and live-cell imaging using Costes randomization.

**Figure S4:**
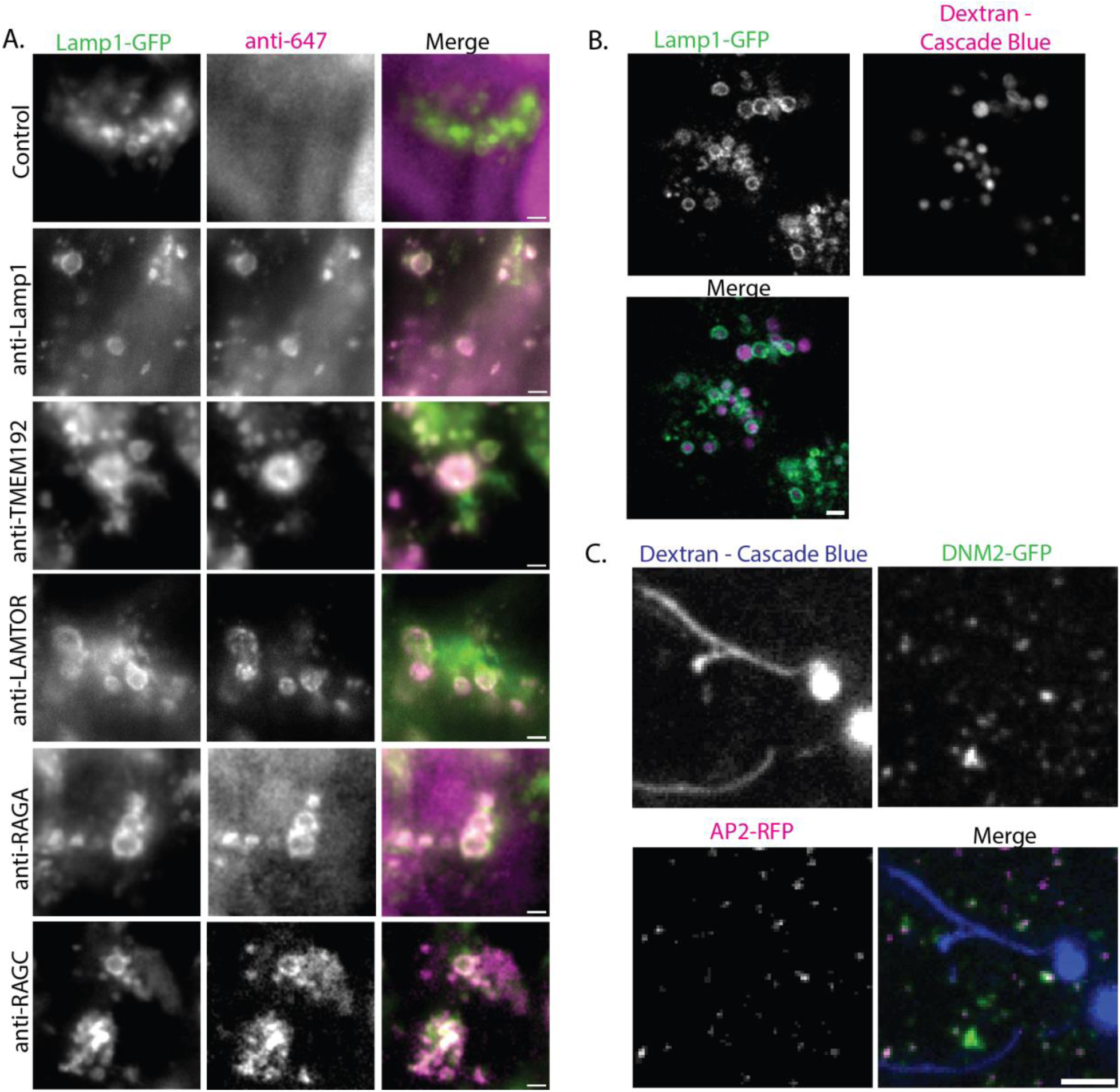
(A) Hi-Lo TIRF imaging of fixed hiPSC LAMP1-GFP expressing cells stained with a variety of primary antibodies to label lysosomal compartments. Scale bar = 2 µm (B) Single z-plane image of LAMP1-GFP expressing hiPSC treated with 10 kDa Dextran-Cascade blue. Scale bar = 2 µm. (C) Single z-plane image of AP2-RFP; DNM2-GFP tagged hiPSCs treated with Dextran-Cascade blue demonstrating that the Dextran can also label lysosome tubules. Scale bar = 3 µm.

